# Miro1 knockout inhibits mouse breast cancer tumorigenesis

**DOI:** 10.1101/2025.05.06.650999

**Authors:** Randi Termos, Alexandra Muskat, Nathaniel Shannon, Margaret McCoy, Ali Termos, Chloe Palmer, Martin C. Chang, Brian Cunniff

## Abstract

Miro1 is a mitochondrial outer membrane protein that regulates mitochondrial and peroxisome trafficking, endoplasmic reticulum (ER) association, and mitophagy. Prior studies suggest a role for Miro1 in cell migration and proliferation in both normal and tumor cells. High Miro1 expression is associated with poor overall survival in breast cancer patients. To investigate the role of Miro1 in breast cancer tumorigenesis and metastasis, we established stable Miro1 knockdown (KD) in MDA-MB-231 human triple-negative breast cancer cells using shRNA. Miro1 KD significantly impaired cell proliferation, migration, and invasion *in vitro*. When implanted into the mammary fat pad of SCID mice, MDA-MB-231 cells formed tumors, whereas Miro1 KD cells showed markedly reduced tumorigenesis. Additionally, we generated a novel transgenic mouse model with inducible, tissue-specific Miro1 deletion in mammary epithelial cells, alongside polyomavirus middle T-antigen (PyVMT) oncogene activation. In this model, wild-type (WT) mice formed tumors at all mammary gland sites, with frequent lung metastases. However, Miro1-deficient mice failed to develop tumors, while heterozygous mice exhibited reduced tumor growth and metastasis. Additionally, these findings identify Miro1 as a key regulator of breast cancer onset and metastatic potential, positioning it as a potential biomarker and therapeutic target.

## Background

Mitochondrial trafficking is mediated by microtubule associated motor proteins, cytoskeletal components, and calcium signaling [1]. Miro1 (mitochondrial Rho GTPase 1) is a key component of this system, embedded in the outer mitochondrial membrane via a C-terminal helix and containing two EF-hand Ca^2+^-binding domains and two GTPase domains [2]. Miro1 serves as an adaptor protein, facilitating mitochondrial transport through interactions with TRAK1/2, which recruit motor proteins to microtubules [3]. Beyond trafficking, Miro1 also regulates mitochondria-ER contacts [4], peroxisome biogenesis [5], [6], and mitophagy [7], [8].

Although the role of Miro1 in cancer development and progression is not completely understood, Miro1 is associated with poor survival, recurrence, and metastasis in multiple cancers [9]. Increased Miro1 expression is associated with gastric cancer progression and is currently being studied as a potential biomarker for early diagnosis and prognosis assessment [10]. Rapidly dividing cancer cells require an increase in ATP production and redox regulation to carry out increased cell division, making mitochondrial dynamics an important component of cancer progression [11]. The recruitment of bioenergetically healthy mitochondria to local sites supports pancreatic tumor cell invasion by providing the required energy to fuel motility and attachment processes [12]. In HaCaT keratinocyte cancer cells, deletion of Miro1 results in inefficient oxidative phosphorylation, reduced ATP at the cell periphery, and significantly altered organization of matrix proteins [13]. Deletion or knockdown of Miro1 in numerous cell types leads to aberrant subcellular mitochondrial positioning [14], [15], [16]. In mouse embryonic fibroblasts (MEFs), deletion of Miro1 leads to perinuclear restricted mitochondria which fail to traffic to the cell periphery [15]. In breast cancer, mitochondria can be transferred between cells, contributing to chemoresistance and disease progression [17]. Mitochondrial anterior positioning was shown to support the directional migration and speed of migration in MDA-MB-231 human breast cancer cells, and modulation of mitochondrial fission (DRP1) and fusion (MFN1/2) proteins slowed their velocity [18]. The migratory and proliferative capacity is also altered in mouse embryonic fibroblasts with Miro1 deletion [14].

Additionally, increased expression of Miro2, which has 60% sequence homology with Miro1, has been shown to be correlated with poor patient survival, recurrence, and metastasis in certain patient cohorts with prostate cancer [19], [20].

Data published on the Human Protein Atlas shows that breast cancer patients harboring high Miro1 expression in their tumor tissue have a poorer survival rate than those with low Miro1 expression [21]. This data suggests that Miro1 expression correlates with patient survival in breast cancer, however, the role of Miro1 in breast cancer is not currently understood.

Here, we generated MDA-MB-231 breast cancer cells with stable Miro1 knockdown to assess tumorigenic properties. Miro1 depletion with shRNA in MDA-MB-231 cells restricted mitochondrial distribution to the perinuclear region that was accompanied by reduced cell proliferation, migration, and invasion. In an orthotopic mouse model, Miro1 KD cells failed to form tumors, while WT cells developed palpable tumors in immunocompromised mice. To validate these findings, we developed a transgenic mouse model with inducible Miro1 deletion in mammary epithelial cells alongside polyomavirus middle T-antigen (PyVmT) driven tumorigenesis. We hypothesized that concurrent Miro1 deletion would reduce tumor formation and metastasis. Miro1 deletion completely prevented tumorigenesis, while heterozygous deletion significantly reduced tumor burden and metastatic potential. These findings highlight Miro1 as a key player in breast cancer progression and provide a novel model for studying mitochondrial dynamics in tumorigenesis.

## Materials and Methods

### Generation of MDA-MB-231 cell lines with Miro1 knockdown

MDA-MB-231 human breast cancer cells were transfected with shCtrl (OmicsLink™ shRNA Expression Clone CSHCTR001-1-mU6; GeneCopoeia, Inc., Rockville, MD) or shMiro1 (OmicsLink™ shRNA Expression Clone, HSH106063-mU6-a; GeneCopoeia, Inc., Rockville, MD) using the GeneJet plasmid miniprep kit (Thermo Scientific #K0502 #K0503) and the X-tremeGENE HP transfection reagent. The plasmid DNA contained an mCherry fluorescent cassette, a puromycin resistance cassette, and an ampicillin resistance gene. Single cells were grown in a 96 well plate to produce colonies, which were selected via fluorescent imaging and puromycin resistance. The initial colonies were then assessed for Miro1 knockdown via qPCR. Clones that had less than 50% Miro1 knockdown were not considered. Two clones, A3 and A4, were propagated for experimental use. The Miro1 knockdown clones and the control cells containing a scrambled plasmid were grown in DMEM/F12 with 10% FBS and 7 µg/ mL puromycin.

### Immunofluorescence staining and analysis

Cells were plated on glass coverslips at 100,000 cells per 6-well dish then fixed for 10 minutes with 4% paraformaldehyde/1x phosphate buffered saline (PBS), then permeabilized with 0.25% permeabilization buffer (triton 100-x in 1x PBS) for 10 minutes. The cells were then blocked in 1.5% BSA/1x PBS at 4°C overnight. The coverslips were incubated with TOMM-20 (1:250, Millipore #MABT166) for 1h, washed in 1x PBS for 40 minutes, incubated with Alexa Fluor 594 phalloidin (1:400, Invitrogen #A12381) and Alexa Fluor 488 (1:500, Invitrogen #A11008), washed in 1x PBS for 40 minutes, then stained with DAPI (0.5μg/mL, ThermoFisher #62248) for 5 minutes. Images were captured using an Eclipse TE-2000E inverted microscope (Nikon) equipped with a 40×/1.3 numerical aperture (NA) Plan Fluor oil-immersion objective. Mitochondrial occupancy (Mito Area/Cell Area) was calculated by isolating regions of interest in the cell leading edge (periphery) and lagging edge and measuring the amount of TOMM20 mitochondrial signal in these regions and that was taken as a percentage of the occupancy in relation to phalloidin in those areas. The mitochondrial form factor was calculated by highlighting the mitochondria and taking their area and perimeter and plugging it into the equation: Form Factor = 0.25 (area/perimeter^2^).

### Wound-healing migration assay

Cells were grown in a 6-well plate; the initial seeding was 3×10^5^ cells in 2 ml DMEM/F12 with 10% FBS, and 7 µg/ ml puromycin for selection. Cells were allowed to grow to approximately 90% confluency for the experiment. A 1000 µl pipette tip was used to gently scrape a line of cells off the plate’s surface to produce a wound. The plates were then loaded into a Lion Heart live-cell imager and were imaged on bright-field setting every hour for 24 hours. The images were then processed in ImageJ software by creating a threshold 8-bit image to select the open wound between the cells and a measurement of the area was taken. This same method was used to produce the highlighted images in Fig.2A. The area for each time-point was calculated to a percentage of closure of the initial wound measurement to account for the inability to make identical wound areas with the initial scraping of cells. The percentages for each time point were averaged across replicate experiments for each cell line and graphed in GraphPad Prism to show percent of wound closure over time with SEM. P-value was calculated using a 2-Way ANOVA test where the source of variation was a row factor for each clone in comparison to the control. N=3

### Transwell invasion assay

Cells were plated in transwell inserts that had a membrane pre-coated with ECM material and 8µm pores (Corning, 354480). FBS-free DMEM/F12 was applied in the insert and DMEM/F12 with 10% FBS in was applied in the bottom chamber. The FBS served as the chemoattractant to stimulate cell invasion. The cells were incubated for 24 hours at 32°C in 5% CO^2^. After the 24-hour incubation, the transwell inserts were removed, fixed in 70% ETOH, stained with 0.1% crystal violet, and washed in distilled water. Cells remaining on the top of the membrane were removed with a sterile cotton tipped applicator prior to staining so their presence would not obstruct the imaging of the invaded cells on the bottom of the membrane. The invaded cells were imaged on a Lion Heart cell-imager on the color bright field setting at 40X magnification. Cell counts were done using BioTek Gen5 Software and those cell numbers were graphed and analyzed in GraphPad Prism. P-values were determined via two-tailed un-paired Student T-tests. N = 3 experimental replicates

### Proliferation assay

Cells were grown in a 6-well plate; the initial seeding was 3×10^4^cell in 2 ml DMEM/F12 with 10% FBS, and 7µg/ ml puromycin for selection. Each following day, one well of the 6-well plate was trypsinized and the number of cells in the solution was counted using a hemocytometer. The measurements were averaged over five repeat experiments and graphed to show cell number over time and SEM in GraphPad prism. P-value was calculated using a 2-Way ANOVA test where the source of variation was a row factor for each clone in comparison to the control. N = 5 experimental replicates

### Xenograft model

At eight weeks of age, female NOD-Scid mice (NOD.CB17-Prkdcscid/NCrCrl, Charles River USA) were injected with MDA-MB-231 human breast cancer cells; control cells with intact Miro1 expression and two clones with approximately 50% Miro1 knockdown were used. Six mice were injected for each cell line. Prepared cells were trypsinized, pelleted, and resuspended in 1X PBS at 1×10^6^cells/ 100 µl PBS, which was the cell suspension volume and concentration injected into each mouse. Mice were anesthetized with isoflurane for the injection procedure. Each mouse received the injection in the right abdominal mammary gland site; as directed by a veterinarian, the injection was made beneath the nipple without an incision. The mice were weighed at the time of injection and their weights were measured every 3 days post-injection. Tumors were measured with calipers every three days after the first palpation. At the point of which the tumorous mice showed signs of discomfort, they were euthanized by CO^2^ and secondary cervical dislocation. Mice that did not form tumors were euthanized at the time of the last tumorous mouse, which was 39 days post-injection. At the time of euthanasia, each mouse was dissected, and all tumor tissue and lung tissue were removed and fixed in 4% paraformaldehyde/ 1X PBS. Tissue samples were later paraffinized and sectioned for staining.

### Generation of Tissue Specific Miro1 KO mice with PyVMT induction

Mice with LoxP sites flanking exon 2 of Miro1 exons (Miro1^FL/FL^) were a kind gift from Dr. Janet Shaw (University of Utah). Mice with the MTB transgene (MMTV-LTR, rtTA) and MIC transgene (Tet-O, PyVMT, IRES, CRE) were acquired from Dr. William Muller at McGill University. MTB and MIC mice were cross bred to produce mice with both transgenes; these mice were then cross bred with Miro1^FL/FL^ mice to produce mice with both the MTB and MIC transgenes, and floxed Miro1 alleles (MTB/ MIC/Miro1^FL/FL^). To generate the control mice, MTB/MIC/Miro1^FL/FL^ mice were bred with MTB/ MIC/Miro1^WT/WT^ mice to breed out the floxed Miro1 alleles and to allow for mixing of the C57BL/6 and FVB/N genomes. From these novel mice, breeding pairs were established to generate control mice with intact Miro1 alleles, heterozygous mice with only one floxed Miro1 allele, and mice with two floxed Miro1 alleles.

### Immunohistochemical tissue staining

Tissue collected from mouse tumors, lungs, and mammary tissue was paraffinized and sectioned by the University of Vermont Medical Center’s pathology laboratory services. The sections were deparaffinized, treated with DAKO antigen retrieval solution and ImmPACT DaB reagent (Vector Laboratories). Sections were stained with Hematoxylin and Eosin (H&E) for visualization of the nucleus and cytoplasm of the cells within the tissue. Sections from the same tissue samples were stained with an antibody for Miro1 (Antibodies Online, ABIN635090) to determine the knockdown of Miro1 in the mammary epithelial tissue of each sample. Additionally, sections of the tissue samples were stained with an antibody to Cre (NOVUS, #NB100-56133) to determine the expression of the transgene. Imaging of the stained slides was done on a Leica VERSA8 Whole Slide Imager.

### Immunofluorescent tissue staining

Tissue collected from mouse tumors and mammary tissue was formalin-fixed, paraffin-embedded, and sectioned by the University of Vermont Medical Center’s pathology laboratory services. The sections were deparaffinized, treated with DAKO antigen retrieval solution. Sections from the same tissue samples were stained with an antibody for PyVMT (NOVUS #NB100-2749) to determine expression of the transgene. The secondary conjugate used was Alexa Fluor™ Plus 647 (Invitrogen #A32733), and DAPI to visualize cell nuclei (0.5 μg/ml, Thermo Fisher Scientific #62248). Images were captured using an Eclipse TE-2000E inverted microscope (Nikon) equipped with a 40×/1.3 numerical aperture (NA) Plan Fluor oil-immersion objective. Images were processed in Image J software as Max-projections of z-stacks with the maximum and minimum brightness and contrast settings fixed across all images for a calibrated assessment of pixel intensity.

### Mammary tumor induction in PyVmT mouse model

After eight weeks of age female mice that had the genotype of interest, as confirmed by PCR genotyping, were put on a diet of doxycycline administered to mice via doxycycline infused food (Isopro 0.6% Doxy (5TWP) Irrad Green per kg., ScottPharma Solutions). Mice were monitored for tumor growth and overall wellness by palpitation of mammary glands, monitoring eating habits and weight, and monitoring for signs of stress or pain. The mice were weighed and palpated every seven days. The number of days up until detectable tumor onset was recorded for comparison between mice with and without Miro1 expression in mammary epithelial cells. Tumors were removed from sacrificed mice seven days after the identification of palpable tumors. Excised tumors were photographed, measured with calipers, and weighed prior to fixation with 4% formaldehyde. In addition to collection of tumor tissue samples from control and experimental mice, normal mammary gland tissue from MTB/ MIC/Miro1^WT/WT^ and MTB/MIC/Miro1^FL/FL^ mice that have not been given doxycycline were collected. The time until tumor onset was graphed as a percentage of tumor-free animals in relation to the time in days post doxycycline administration.

Tissue samples taken from sacrificed mice were fixed, embedded in paraffin and sectioned onto slides for Hematoxylin and Eosin (H&E) staining. With guidance from a diagnostic pathologist with experience in human and mouse histology (Chang), we analyzed the H&E-stained slides and recorded changes in histological architecture, hyperplasia, anaplasia, necrosis, and the presence of metastasized tumor cells to the lungs. Expression of PyVMT, CRE and Miro1 were determined by IHC staining to confirm all components of the model were functioning. Graphing of tumor scoring, percentage of mice with lung metastases, and percentage of mice with tumors was done in GraphPad Prism. P-values for scored tumor analysis were generated using two-tailed unpaired T-tests comparing the HET scores to the WT scores, and the FL/FL scores to the WT scores.

### qPCR for Miro1 expression in MDA-MB-231 cells

RNA was isolated and purified from MDA-MB-231 cells using Qiagen’s RNeasy mini kit (Qiagen ID. 74104). The protocol was carried out as per the manufacturer’s instructions. PCR amplification of the RNA product was carried out using a Qiagen RT^2^ first strand kit (Qiagen ID. 330401) in preparation for quantitative PCR; 500 ng RNA was used for each reaction. The cDNA product was diluted 1:4 for the qPCR reactions. RT^2^ SYBR Green Fluor qPCR Mastermix (Qiagen ID. 330513) was added to the diluted product. Each qPCR reaction set included a negative control reaction without a template. qPCR was preformed using PrimeTime qPCR primers from Integrated DNA Technologies (IDT, Madison, WI, USA). The qPCR reactions were run with the following thermocycler conditions: (95°C X 10 minutes, 95°C X 15 seconds, 60°C X 1 minute) X40 cycles.

Relative gene expression levels were calculated using the 2^-ΔΔCt method. ΔCt was determined by subtracting the Ct value of the endogenous control gene (ACTb) from the Ct value of the target gene for each sample. ΔΔCt values were then calculated relative to a designated calibrator sample (MDA-MB-231 shCtrl cell RNA). Fold changes in gene expression were determined as 2^-ΔΔCt. Data was graphed using GraphPad Prism. n=3 per group.

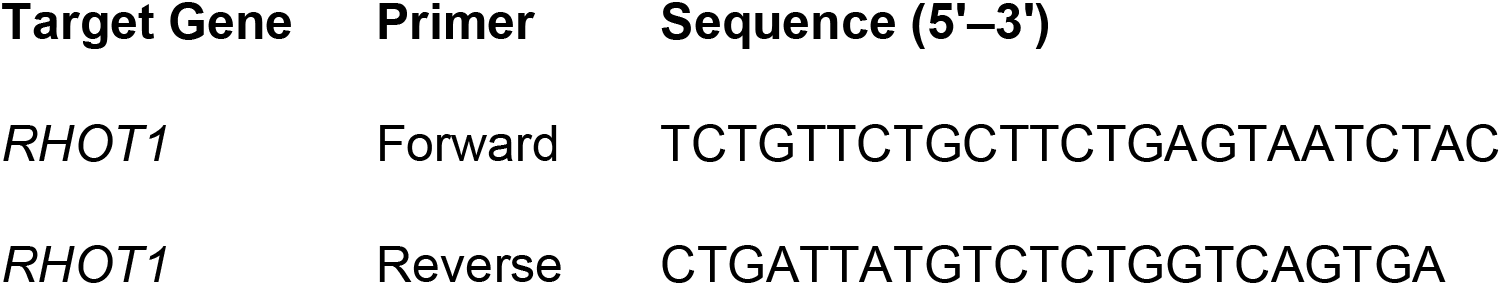

### Genotyping of transgenic mice

Ear tissue clippings were collected and subjected to DNA extraction using the Applied Biosystems DNA Extract All Reagents Kit (Cat# 4403319) according to the manufacturer’s instructions. Samples were either used immediately or stored at –20°C until use. PCR was preformed to amplify the gene of interest. One microliter of the extracted DNA dilution was used in each PCR reaction. Positive controls and a negative (no template) control were included. Thermal cycling conditions were optimized for each primer set. PCR products were resolved on 2% agarose gels prepared with ethidium bromide. Gels were run at 125 V for 50 minutes and visualized under UV light. Representative expected amplicon sizes were approximately 500 bp (MTB), 800 bp (Cre), and 150 bp and 100 bp (Miro1 floxed alleles). Primers were custom-designed and synthesized by Integrated DNA Technologies (IDT, Madison, WI, USA).

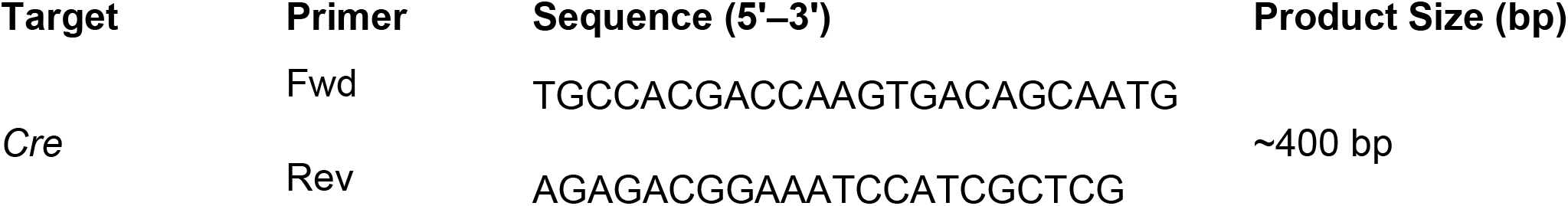
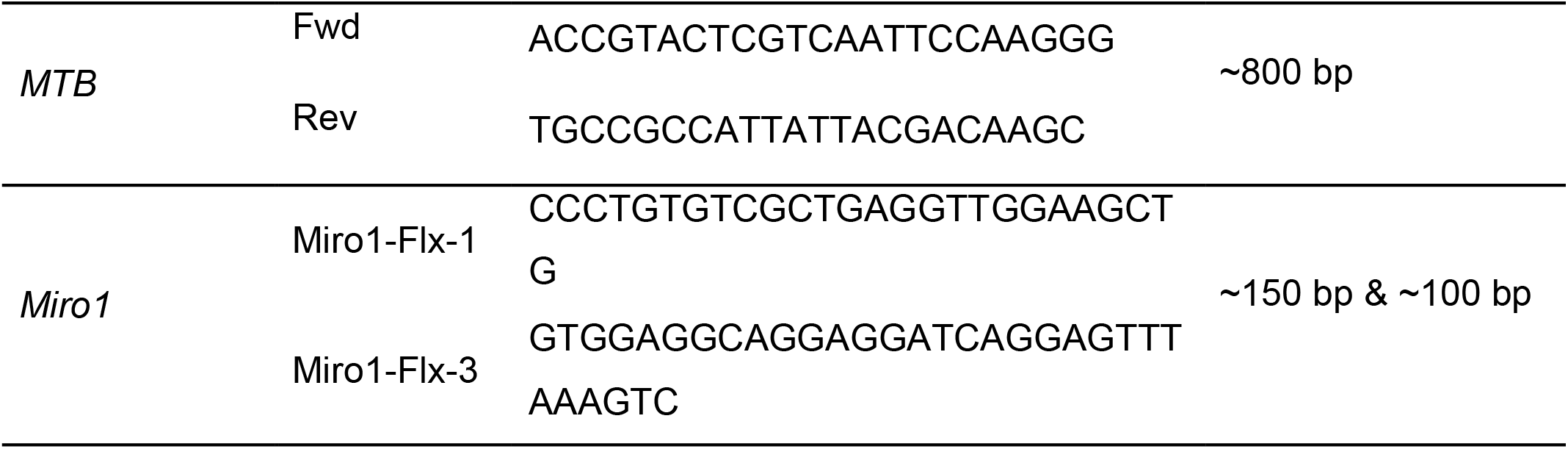

Genotype was assessed by presence or absence of a band on an agarose gel. PCR products were further validated through sequencing at genomics core at The University of Vermont. The resulting sequences were analyzed in FinchTV software, then input into the National Center for Biotechnology Information’s (NCBI) Basic Local Alignment Search Tool (BLAST); the resulting sequences aligned with similar reference sequences for the target genes.

## Results

### Stable knockdown of Miro1 leads to perinuclear restricted mitochondria, and reduced cell proliferation, migration and invasion

To evaluate the role of Miro1 in human breast cancer cells, we generated MDA-MB-231 cells with stable knockdown of Miro1. Miro1 KD clones A3 and A4 both exhibited approximately 50% knockdown in comparison to control cells as determined by RT-qPCR (Fig 1B &C). Depletion of Miro1 resulted in mitochondria restricted to the perinuclear region of the cell (Fig. 1A &D), as has been observed in other cell types [14], [15], [16]. Additionally, mitochondrial networking was significantly increased in Miro1 knockdown cells (Fig. 1F).

**Figure 1.**
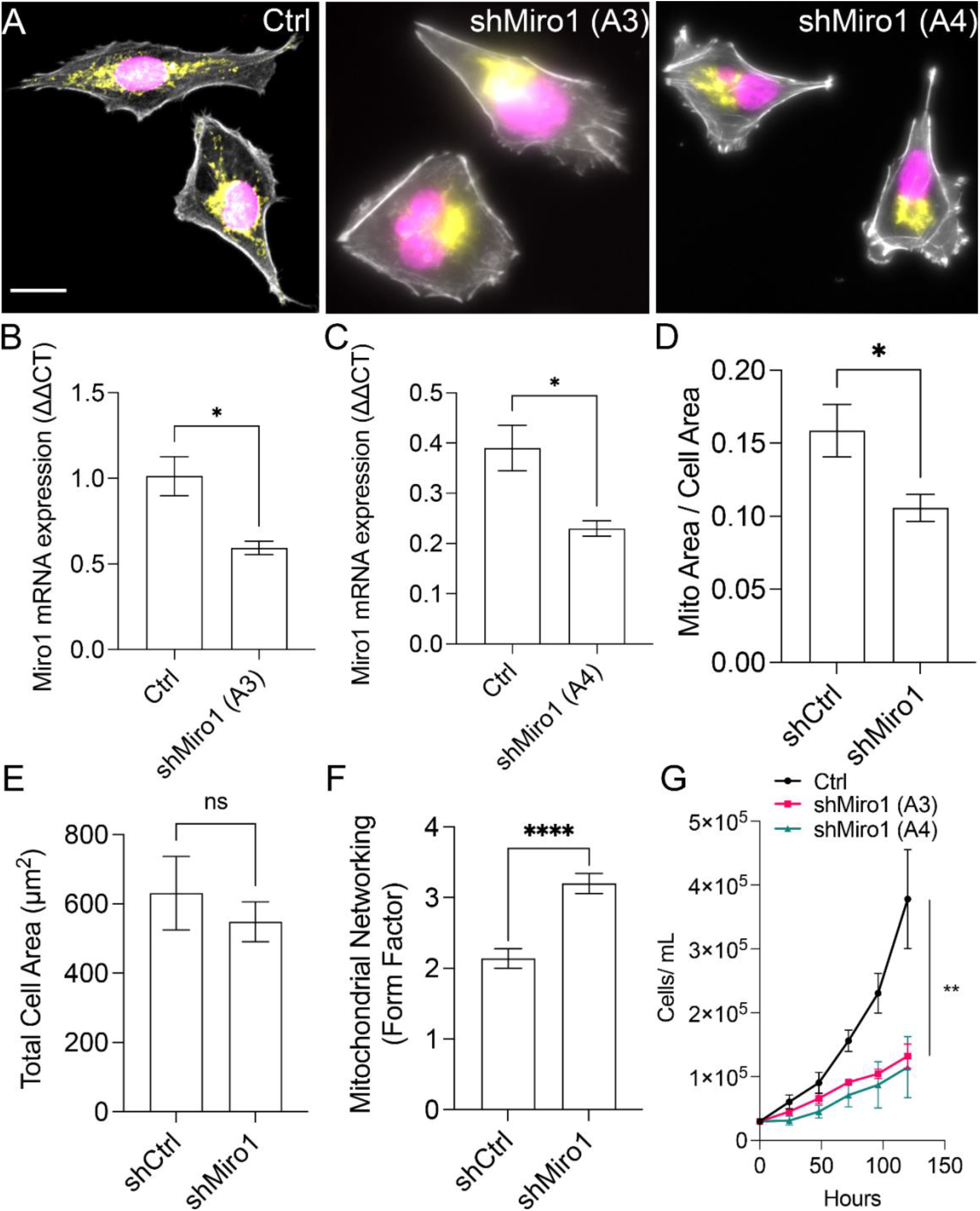
Miro1 knockdown restricts mitochondria to perinuclear regions and inhibits cell migration of MDA-MB-231 breast cancer cells. **A)** MDA-MB-231 cells were transfected with control or Miro1 shRNA prior to fixation on glass slides and immunofluorescence staining. Purple = DAPI/nucleus, Yellow = Anti-TOMM20 antibody/ mitochondria and Gray = Phalloidin/Actin. **B and C**) Miro1 mRNA expression in stable MDA-MB-231 Miro1 shRNA clones measure by RT-qPCR (n = 3, * p < 0.05, Students t-test). **D**) Quantification of mitochondria area relative to cell area (n = 20 cells/group, * p < 0.05, Students t-test). **E**) Total cell area is unchanged in shCtrl and shMiro1 cells (n = 20 cells/group). **F**) Mitochondrial networking is increased in shMiro1 cells (n = 20 cells/group, **** p < 0.001, Students t -test) **G**) Stable knockdown of Miro1 in MDA-MB-231 clones’ knockdown significantly reduces cell proliferation (n = 5, ** p < 0.01, One-way ANOVA with Tukey’s Post Test).

Miro1 KD cells exhibited a significant decrease in cell proliferation (Fig. 1G) and significantly decreased cell migration in wound healing assays compared to control cells (Fig. 2A, B &C). Miro1 KD cells also had reduced ability to invade in transwell invasion assays compared to control cells (Fig. 2C &D). Together, these data provide evidence for Miro1 supporting MDA-MB-231 breast cancer cell growth, migration, and invasion.

**Figure 2.**
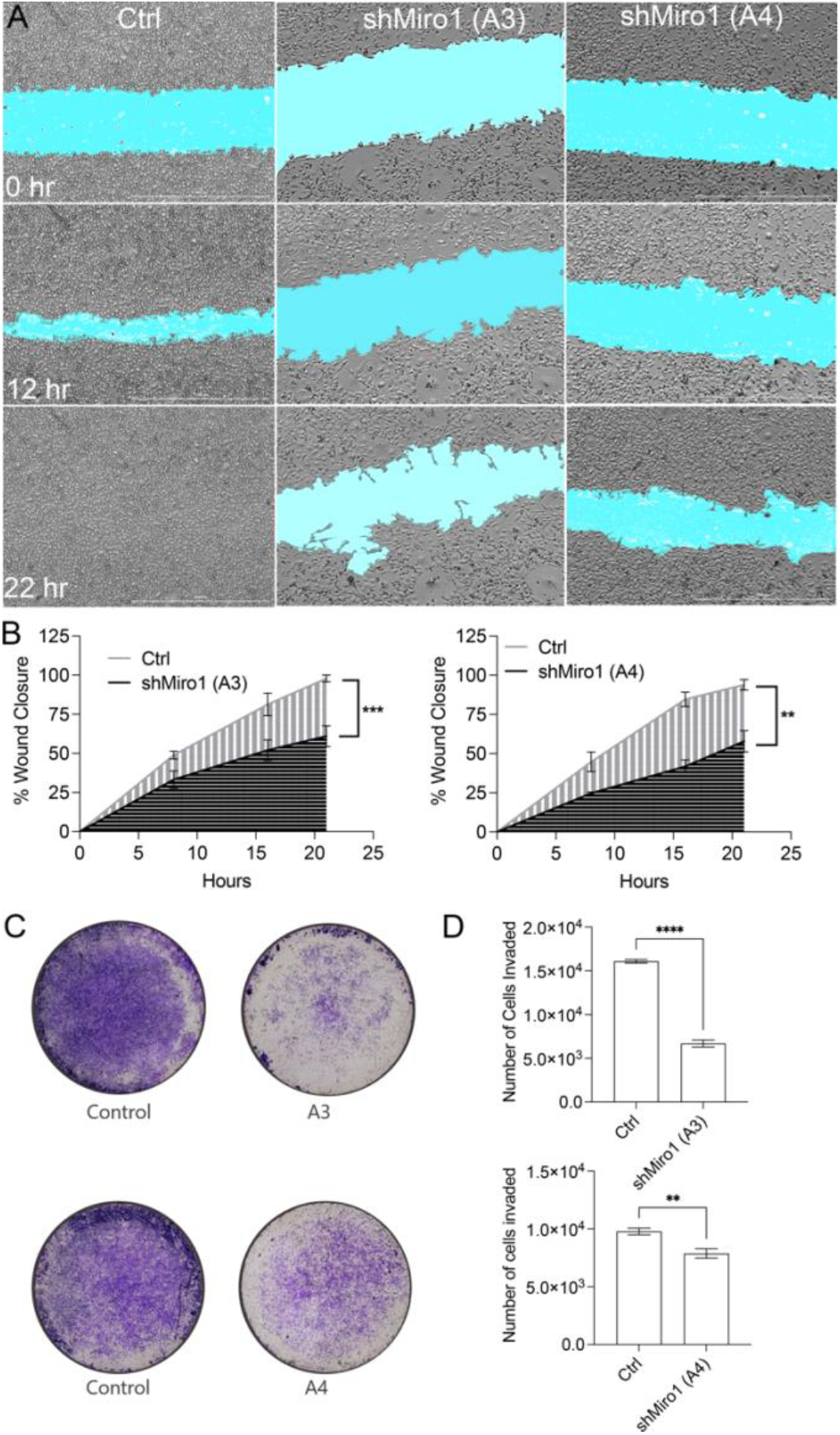
Miro1 knockdown reduces MDA-MB-231 cell migration, proliferation and invasion. (**A&B**) MDA-MB-231 clones with Miro1 knockdown had significantly reduced ability to close the gap in wound closure migration assays. Open wound area highlighted in cyan. (**C&D)** MDA-MB-231 cells with Miro1 knockdown had significantly reduced ability to invade through a membrane in transwell invasion assays. Invaded cells were stained with 0.1% crystal violet.

### Stable knockdown of Miro1 prevents MDA-MB-231 tumor cell growth in an orthotopic mouse model

To further evaluate tumorigenic properties of MDA-MB-231 cells with stable Miro1 KD, we conducted orthotopic tumor implantations of MDA-MB-231 Miro1 KD cells into the mammary fat pad of immunocompromised mice. Six female NOD-SCID mice per group (Control, shMiro1 A3 clone, and shMiro1 A4 clone) were injected with 1×10^6^ cells into the mammary fat pad and tumor burden was monitored daily via palpitation (Fig. 3A). Tumors were measured with calipers two times per week until the time of euthanasia and were palpable in the control mice after an average of 13 days. Nine out of twelve (75%) control mice formed mammary tumors at the injection site between days 8-26 (Fig. 3A). Two out of twelve (17%) shMiro1 clone A4 mice formed mammary tumors at the injection site at an average of 14.5 days. All mice that received injections of the shMiro1 clone A3 failed to form palpable tumors (Fig. 3A). To visualize tumor burden, tissue was collected for Immunohistochemical analysis (Fig. 3B). Animals injected with control cells formed large, dense tumors with necrotic cores while mammary tissue from mice harboring Miro1 KD cells showed normal histology (Fig. 3B). Tumor burden was scored from blinded slides by an anatomical pathologist using a standardized scoring criterion (Supplementary Fig. S1). Ctrl animals had a significantly higher tumor score compared to shMiro1 tissue (Fig. 3C). Together, stable KD of Miro1 inhibits tumor growth following orthotopic injection into the mammary fat pad.

**Figure 3.**
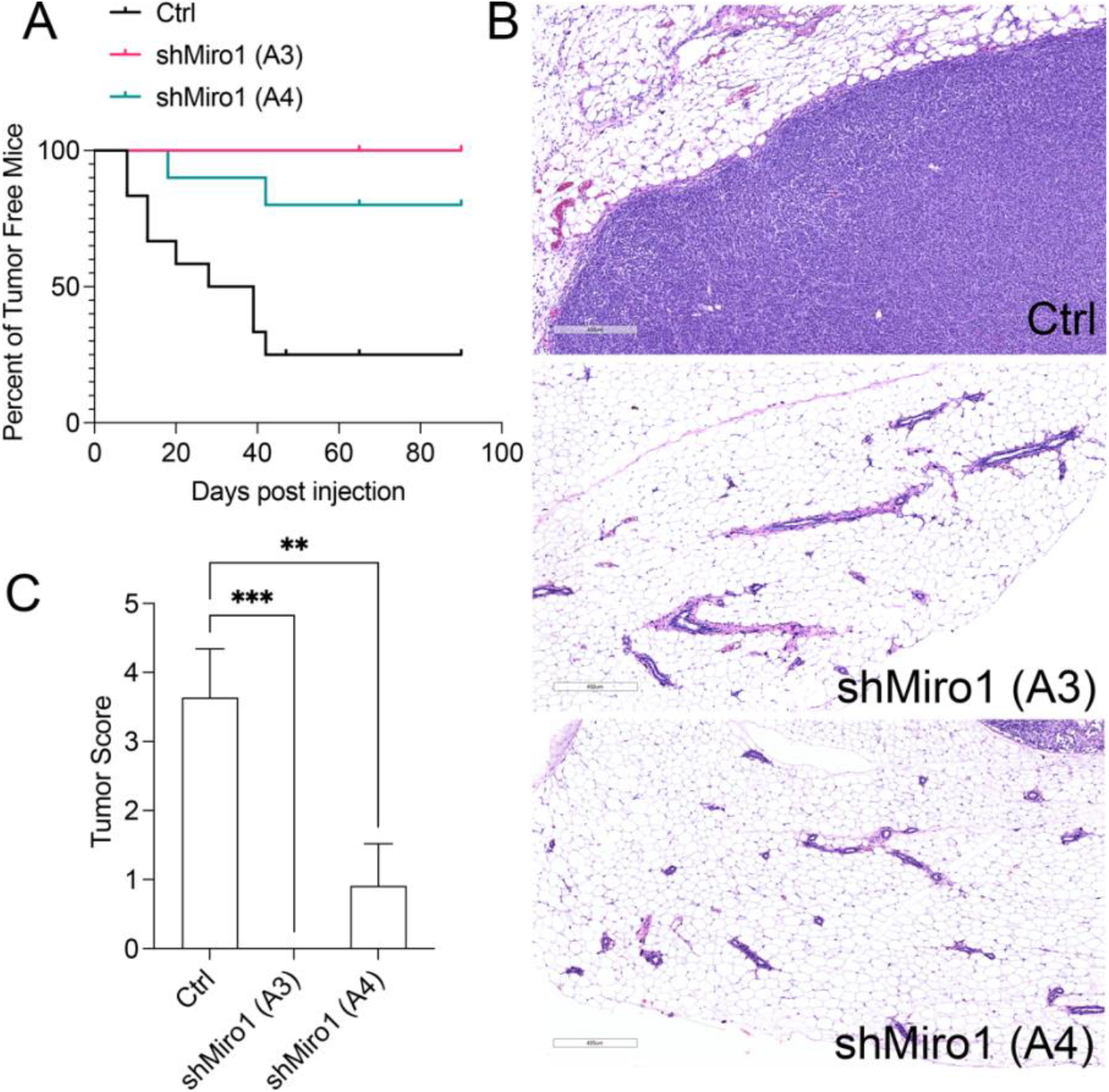
Miro1 Knockdown reduces MDA-MB-231 tumor growth *in vivo*. (**A**) Percentage of tumor-free mice over days post-orthotopic injection of MDA-MB-231 clones with and without Miro1 knockdown. 75% of control mice formed mammary tumors at the injection site between days 8-26. Only 17% of mice injected with the Miro1 knockdown clone, A4, formed mammary tumors at the injection site between days 11-18. All mice that received injections of the Miro1 knockdown clone, A3, failed to form palpable tumors. (**B**) Scoring of tumors (Score 1-5) post excision from mice during necropsy. Scoring was done using tissue slides stained with hematoxylin and eosin (H&E). (**C**) Scoring of mammary epithelial tumor tissue based on our assessment of histological features such as loss of normal architecture, cellular morphology, and stromal invasion (Supplementary figure 1).

### Generation and characterization of transgenic Miro1 KO mouse model

To investigate breast cancer tumor growth and metastasis in an advanced mouse model, we generated a novel mouse model that has concurrent polyomavirus middle T antigen (PyVMT) mammary epithelial tumor initiation and deletion of Miro1 (Fig. 4A). The model utilizes the TetO-PyVmT-IRES-CRE recombinase (MIC) transgene, which links the oncogenic PyVMT protein with CRE recombinase in a bicistronic configuration. Site specificity is driven by a mammary epithelial-specific MMTV promoter, which is linked to the rtTA gene (MTB) (Fig. 4A). The MTB/ MIC mouse model is a well characterized inducible system designed to study mammary tumorigenesis with temporal and spatial control over transgene expression [22], [23].

**Fig. 4.**
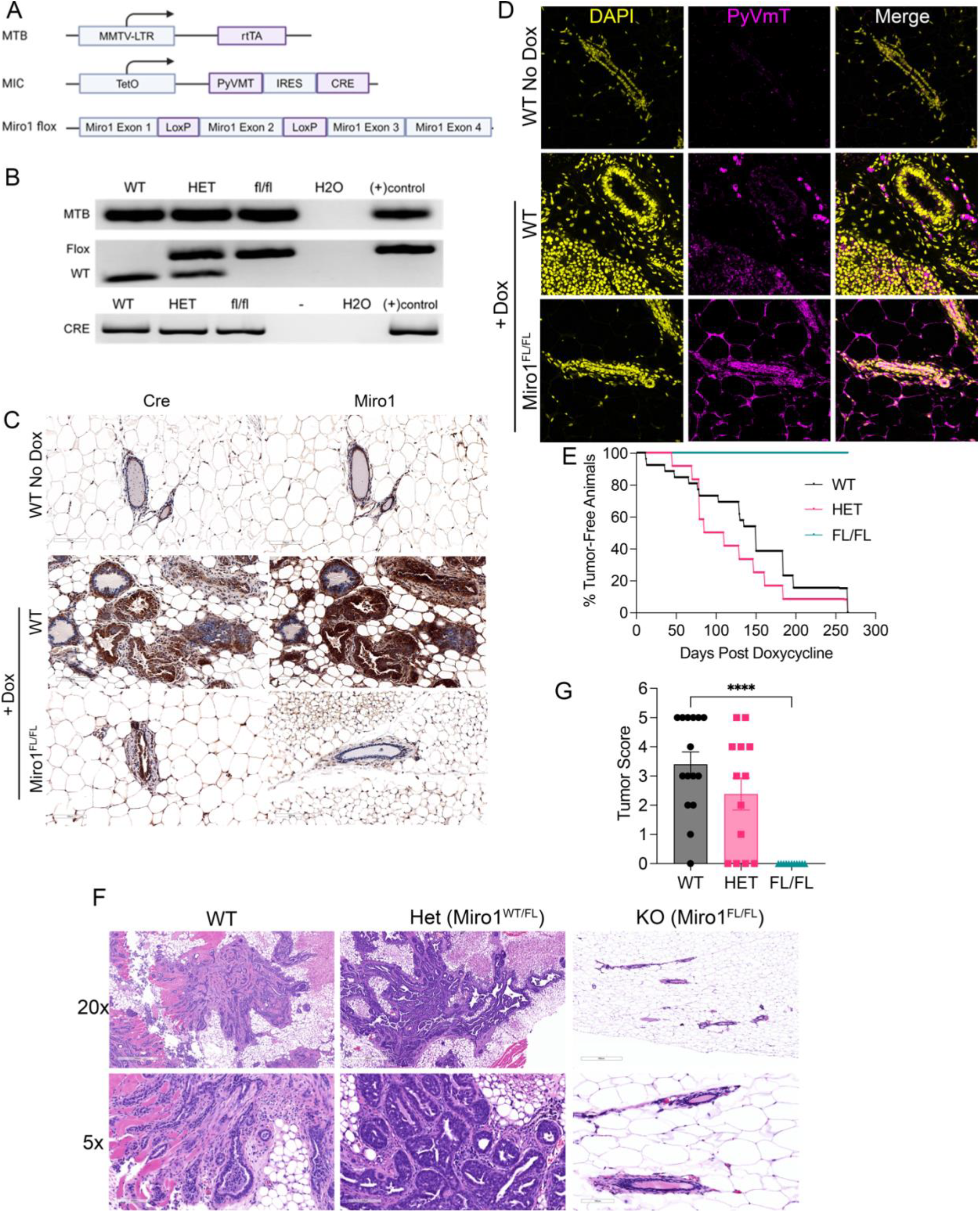
Deletion of Miro1 from mammary epithelial cells blunts tumor formation following PyVmT expression. **(A)** Generation of the transgenic mouse model: Control mice have middle T-antigen activation and the experimental mouse model has concurrent Miro1 deletion and middle T-antigen activation. Cross breeding of the MIC mouse and the MTB mouse models with the Miro1 ^fl^/fl murine model generated the final desired experimental mouse (MIC/ MTB/ Miro1 ^fl^/fl). Activation of the mammary tumor virus long terminal repeat (MMTV-LTR) promoter in the presence of hormones results in reverse tetracycline trans activator (rtTA) production. In the presence of tetracycline (doxycycline) and rtTA, the tetracycline-inducible operator (TetO) is activated, leading to the simultaneous production of Polyomavirus middle T antigen (PyVMT) and CRE recombinase (CRE). An internal ribosome entry sequence (IRES) ensures simultaneous gene expression. **(B)** Presence of the transgenes were confirmed by genotyping an ear-clipping sample from each mouse used in the experiment. **(C)** IHC staining of Cre (where Cre is in brown and cell nuclei are in blue) confirmed expression in both WT and KO mice. IHC staining confirmed expression of Miro1 (where Miro1 is in brown and cell nuclei are in blue) in WT mice and lack of expression in KO mice. **(D)** IF staining of PyVMT (where PyVMT is yellow, and cell nuclei are pink) confirmed its expression in both the WT and KO mice. **(E)** percentage of tumor-free mice over time in days post-doxycycline administration. **(F)** Hematoxylin and eosin (H&E) staining of mammary epithelial tissue (where purple is cell nuclei and pink is cytoplasm) collected from wild type, ^fl^/fl (Miro1 knockout), and heterozygous mice. **(G)** Scoring of tumors (1-5) removed from mice during necropsy using one representative tumor per mouse. Scoring of mammary epithelial tumor tissue based on our assessment of histological features such as loss of normal architecture, cellular morphology, and stromal invasion. (Supplementary Figure 1).

Upon administration of doxycycline (Dox) via infused feed pellets, rtTA forms a complex with Dox and activates Tet-O, driving concomitant expression of PyVMT and CRE. PyVMT expression leads to mammary tumor development and progression that closely simulates human breast cancer [22], [23]. MTB/MIC mice were cross bred with mice that harbor LoxP sites flanking exon 2 of Miro1 as described by Ngyuen et al. [24]. This resulted in the generation of WT mice (MTB/MIC^WT^/^WT^), heterozygous mice (MTB/MIC^WT^/^FL^), and Miro1-KO mice (MTB/MIC^FL/FL^). The Dox inducible system enables the initiation of tumor formation at a specified time (at the age of sexual maturity), and CRE recombinase mediates simultaneous Miro1 deletion in mammary epithelial cells [22], [23]. The presence of the transgenes and Miro1 LoxP containing alleles in mouse pups was confirmed through genotyping prior to the study (Fig. 4B). Mammary epithelial tissues were collected from WT and Miro1^FL/FL^ mice to assess expression of Cre recombinase, Miro1, and PyVmT by Immunohistochemistry and Immunofluorescence (Fig 4. C & D). WT mice that were not given Dox feed show no expression of Cre or PyVmT (Fig. 4 C & D). Cre and PyVmT expression was present in mammary epithelial cells from WT and Miro1^FL/FL^ mice upon administration of Dox food (Fig. 4 C & D). Cre and PyVmT expression was seen in a WT mouse after 12 days of doxycycline containing diet. Miro1 expression was present at low levels in WT mice without Dox food while expression is significantly higher in WT animals fed Dox and harbor visible tumor formation (Fig. 4C). Miro1 staining is absent from epithelial cells of Miro1^FL/FL^ mice in the presence of a Dox containing diet (Fig. 4C).

Our hypothesis was that Miro1 deletion would slow tumorigenesis and inhibit metastasis given Miro1’s role in cell migration [14], [25], [26]. However, all mice with Miro1 deletion failed to form tumors (Fig. 4E). 95% of control mice formed mammary tumors and 71% of those mice had metastasis to the lungs (Supplementary Fig. S2). Heterozygous Miro1 deletion resulted in the formation of tumors equal to WT mice (Fig. 4E) with decreased presence of lung metastasis (Supplementary Fig. S2). This data shows that loss of Miro1 inhibits tumorigenesis in mice with PyVMT initiated cell transformation and decreases metastasis to the lungs.

Tissue from all the mice was collected from each of the mammary gland sites as well as the lungs (Fig. 4F). The tissue samples were stained with hematoxylin and eosin (H&E) to assess tumor morphology and presence of lung metastases (Fig. 4F); the tumors were scored from blinded slides by an anatomical pathologist using a standardized scoring criteria (Supplementary Fig. S1) to assess tumor progression across the genotypes (Fig. 4G). Miro1 WT and Het animals had similar tumor burden scores while no tumors were detectable in Miro1 KO tissues (Fig. 4G). High scoring tumor tissue from control animals showed invasive adenocarcinoma with confluent expansile growth patterns. Regions of squamous metaplasia and keratinization were present in some samples. The tumor cells were high grade, with markedly increased nuclear-to-cytoplasmic ratio. Miro1 knockout mammary epithelial tissue reportedly showed normal-appearing fat tissue and benign glandular elements having ductal arrangement.

## Discussion

Our work demonstrates a critical role for Miro1 in breast cancer tumorigenesis, proliferation, and metastasis. Using our novel transgenic mouse model that has Miro1 deletion in mammary epithelial cells, and orthotopic xenografts of MDA-MB-231 triple-negative breast cancer cells with Miro1 knockdown, we found that reduced Miro1 expression significantly blunted tumor formation in both mouse models and decreases metastatic potential in functional assays *in vitro*. These findings align with our ongoing research investigating the mechanism linking Miro1-mediated mitochondrial positioning to pathways involved in cell migration [14], [27], signaling [26], tumorigenesis, and metastasis.

The perinuclear restriction of mitochondria observed in MDA-MB-231 cells with stable Miro1 knockdown supports previous findings that Miro1 is essential for proper mitochondrial trafficking [14], [20]. The inhibition of mitochondrial distribution is associated with reduced cell proliferation, migration, and invasion although we cannot differentiate whether the effects are related to Miro1 deletion or the perinuclear clustering of mitochondria in this model and this point requires further investigation.

These *in vitro* phenotypes align with *in vivo* results, where cells with reduced Miro1 expression exhibited a markedly reduced ability to form tumors in orthotopic models. The increased mitochondrial networking observed in our knockdown cells may be a contributing factor to impaired trafficking, as mitochondrial fission has been shown to support tumor cell migration [28]. Published work has shown that in cells with inhibited mitochondrial fission protein expression, mitochondria are larger, and their networking or fusion impedes their trafficking to the leading edge [18]. These cells migrate at a lower velocity, which suggests that altered mitochondrial fission and fusion may disrupt cellular processes that are essential for metastasis.

Additionally, published work shows that Miro1 regulated the redistribution of mitochondria to areas of high ATP demand, particularly during migration [14]. Cancer cells may rely on localized ATP outputs to sustain cytoskeletal remodeling and membrane protrusions during invasion; published work shows that metastatic progression of pancreatic cancer cells can be driven by changes in ATP/ADP and ATP/AMP ratios [29], [30]. Recent work underscores this by showing that distinct populations of mitochondria are found at filopodia, and this supports cell migration phenotypes [27]. The perinuclear clustering of mitochondria in our Miro1-deficient cells may create an energy deficit at the cell periphery, thus impairing motility and invasion. It has been shown in MEF’s with Miro1 deletion, the mitochondria were bioenergetically healthy, and their outputs were unchanged, however the localization of those outputs appears to be sequestered to the local sites of the mitochondria [15]. If ATP and ROS are unable to reach the periphery of the cell, signaling cascades affecting attachment, migration, and invasion could be affected in metastatic cancer cells.

The reduced metastatic potential observed in our transgenic Miro1 heterozygous mice may highlight a dose-dependent role of Miro1 in breast cancer. While tumors formed in these mice, their ability to invade distant sites, such as the lungs, was significantly impaired. This implies Miro1-mediated mitochondrial positioning is utilized not only in primary tumor formation, but also the invasive and metastatic potential for cancer progression. These findings are consistent with prior studies linking Miro1 expression to poor patient survival and increased metastasis in other cancer types, including gastric [10] and prostate cancers [19]. Additionally, this data compliments our *in vitro* assays in which Miro1 knockdown resulted in reduced migration, invasion, and proliferative phenotypes.

While the mechanism of Miro1 utilization in tumorigenesis and metastasis requires further research, our current data positions it as a potential biomarker for breast cancer prognosis and a potential therapeutic target. The complete inhibition of tumor formation in our Miro1 knockout models suggests that strategies to inhibit Miro1 expression or function could effectively suppress tumor progression. Strategies to target Miro1 pharmacologically have been explored previously in models of Parkinson’s disease. The Miro1 reducer compound identified in these studies targets Miro1 for proteasomal degradation in the presence of a mitochondrial membrane potential uncoupler [31]. It is possible that inherent differences in membrane potential of tumor cells may limit or enhance the activity of this compound, warranting further investigation.

While our study provides compelling evidence for the role of Miro1-mediated mitochondrial positioning in breast cancer, several questions remain. The specific molecular mechanisms by which Miro1 deletion alters signaling pathways in cancer cells require further elucidation. Emerging research indicates that Miro1 significantly influences various cellular signaling pathways beyond its established role in mitochondrial transport. Studies by Shannon et. al identified significant changes in the expression of proteins associated with cell cycle progression and the MAP kinase pathway in MEFs, which identifies potential signaling pathways altered by Miro1 loss that would be of interest to investigate in breast cancer models [16]. Disruption of Miro1 has been also associated with altered PINK1-Parkin mitophagy pathways, which are critical for turnover of damaged mitochondria [32]. The inability of Miro1-deficient cells to efficiently clear damaged mitochondria may lead to increased oxidative stress and altered apoptosis signaling [33], offering another potential mechanism to investigate by which Miro1 inhibition may suppress tumorigenesis. Considering these findings, future research efforts may focus on addressing whether the clustering of mitochondria around the nucleus in Miro1 depleted cells contributes to oxidative stress or oxidative damage at the nucleus, thus initiating DNA damage controls and cell senescence.

Future directions will include the generation of MDA-MB-231 cell lines with over-expressed Miro1 to investigate the growth, migration, and invasion phenotypes further. Exploring the link between Miro1 inhibition and other proteins of interest in breast cancer tumorigenesis and metastasis will further our understanding of their cooperative or compensatory roles in cancer progression. In preliminary studies we have uncovered altered phosphorylation events and altered protein expression involving key signaling proteins. The mechanism by which these pathways are altered requires further research. Additionally, we aim to generate transgenic mice with non-concomitant deletion of Miro1 so we can investigate the effects of its knockout after tumor growth has been initiated. Our current transgenic model provides us with heterozygous deletion of Miro1, in which the mice display decreased metastasis to the lungs. We aim to investigate to what degree dose-dependent Miro1 expression results in decreased metastatic phenotypes.

In conclusion, our findings establish Miro1 as a critical regulator of mitochondrial dynamics, essential for breast cancer tumorigenesis and metastasis. These results build the foundation for further investigation of Miro1 utilization in breast cancer. By elucidating the molecular mechanisms through which Miro1 mediated mitochondrial positioning influences breast cancer tumorigenesis and metastasis, we may uncover novel strategies for targeting aggressive breast cancer subtypes.

## Declarations

### Ethical Approval and Consent to participate

Not Applicable

### Consent for publication

Not Applicable

### Availability of supporting data

Primary data will be made available upon request

## Authors’ contributions

RT, MM, AT, CP, AM and MCC acquired and analyzed data. BC and RT wrote the initial manuscript draft. All authors contributed to editing and reviewing. BC and MCC obtained funding and BC provided management of the project

## Acknowledgements

We thank Dr. David Seward, University of Vermont Department of Pathology and Laboratory Medicine for generous use of his Lionheart Imaging System. Imaging work was performed at the Microscopy Imaging Center at the University of Vermont (RRID# SCR_018821). Confocal microscopy was performed on a Nikon A1R-ER point scanning confocal supported by NIH award number 1S10OD025030-01 from the Office of Research Infrastructure Programs. A Larner College of Medicine Shared Instrumentation Award was used to purchase the Leica-Aperio VERSA whole slide imaging system. We thank Nicole Bouffard for technical assistance with tissue processing and imaging. We thank Dr. William Muller (McGill University) and Dr. Lewis Chodosh (University of Pennsylvania) for providing the MTB/MIC mice.

## Ethics

All protocols used in animal experiments were approved by the University of Vermont College of Medicine Institutional Animal Care and Use Committee (IACUC).

**Supplemental Figure 1:**
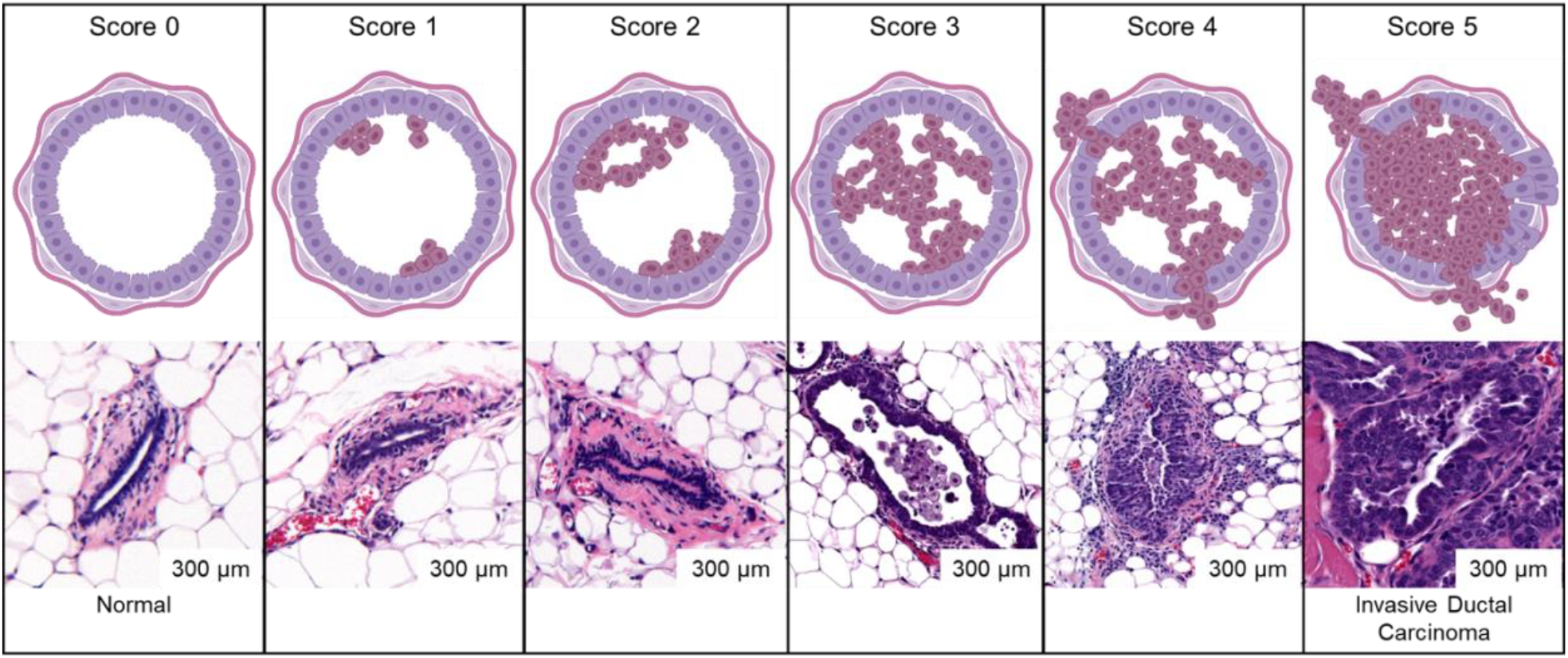
Scoring chart of H&E-stained mammary duct tissue. (Top row) Schematic representation of stepwise neoplastic transformation of mammary ducts. Scoring begins at 0 with a normal ductal structure, followed by hyperplasia, atypical hyperplasia, ductal carcinoma in situ (DCIS), and culminates in invasive carcinoma characterized by cellular overgrowth and breaching of the basement membrane. (Bottom row) Representative H&E-stained histological sections of mouse mammary ducts corresponding to each score. Normal ducts exhibit a single layer of epithelial cells outlining the duct, while increased scoring of tissue shows increased cellular proliferation and thickening, architectural distortion, nuclear pleomorphism, and eventual stromal invasion. Scale bars = 300 µm.

**Supplemental Figure 2:**
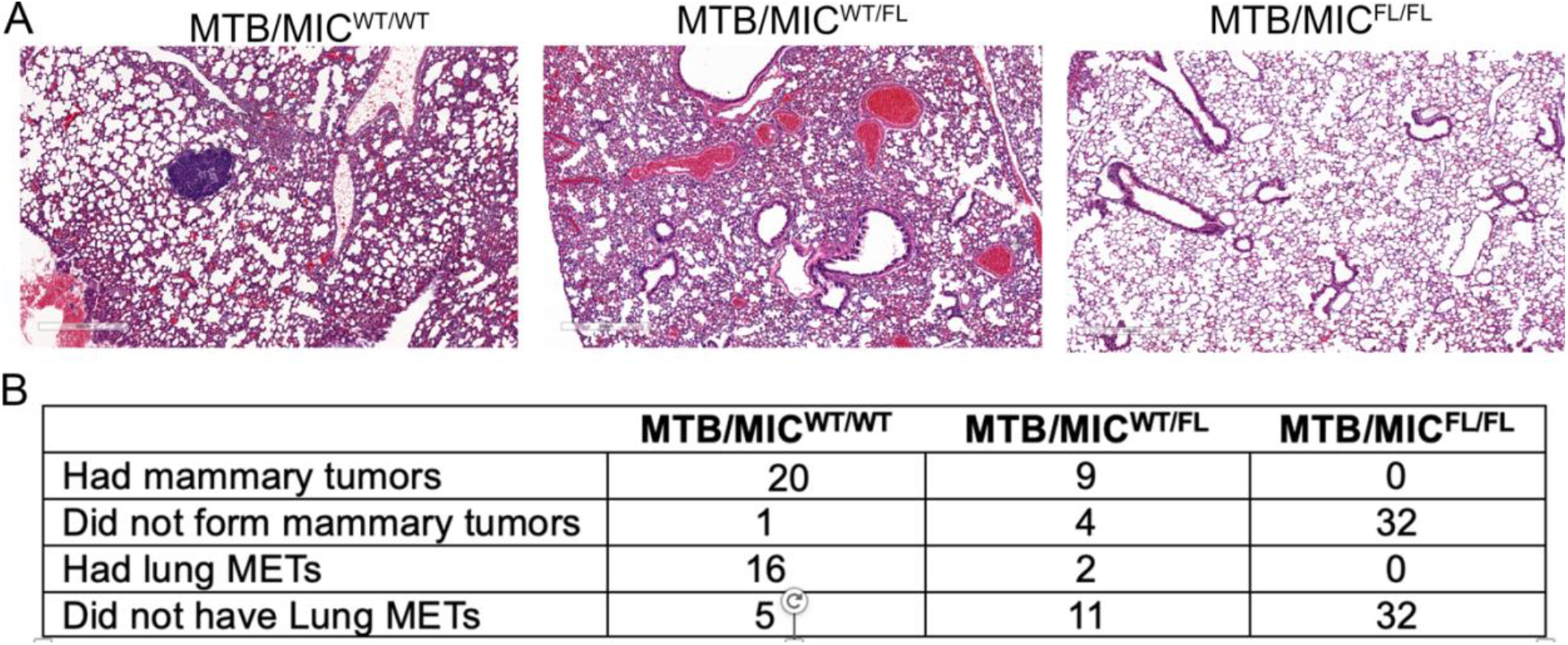
Miro1 Knockout and heterozygous expression in mammary tissue reduces metastatic potential. (**A**) H&E-stained lung tissue from indicated mice (**B**) Table showing the number of animals with and without primary mammary tumor formation and with and without metastasis to the lungs.

